# Bioconversion of pomegranate residues into biofuels and bioactive lipids

**DOI:** 10.1101/2021.04.27.441664

**Authors:** Marianna Dourou, Christina N. Economou, Lida Aggeli, Miroslav Janák, Gabriela Valdés, Nefeli Elezi, Dimitrios Kakavas, Theodore Papageorgiou, Dimitrios V. Vayenas, Milan Certik, George Aggelis

## Abstract

Pomegranate residues (PRs) (i.e. the solid residues remaining after juice extraction), generated currently in abundance in Greece, contain a variety of carbon sources and therefore can be regarded as a potential feedstock for chemical and biotechnological processes rather than as waste materials. In the current project, the polysaccharides contained in PRs were extracted and hydrolyzed in a one-step process without the use of chemical reagents and the resulting broth was used as substrate in biotechnological applications, including ethanol and single cell oil (SCO) production. The yeasts *Meyerozyma guilliermondii*, *Scheffersomyces coipomoensis*, *Sugiyamaella paludigena* and especially *Saccharomyces cerevisiae*, were able to efficiently convert PR derived reducing sugars into bioethanol. Ethanol production under anaerobic conditions ranged from 3.6 to 12.5 g/L. In addition, the oleaginous yeasts *Lipomyces lipofer* and *Yarrowia lipolytica* as well as *M. guilliermondii*, *S. coipomoensis* and *S. paludigena* were tested for their ability to accumulate lipids suitable as feedstock for biodiesel production. Lipids were accumulated at concentrations up to 18% and were rich in palmitic acid (C16:0) and oleic acid (C18:1). Finally, the oleaginous fungus *Cunnichamella echinulata* was cultivated on PR based solid substrates for γ-linolenic acid (GLA) production. The fermented bio-products (i.e. fermented substrate plus fungal mycelia) contained up to 4.8 mg GLA/g of dry weight. Phenolic removal (up to 30%) was achieved by several of the above mentioned microorganisms, including *C. echinulata*, *L. lipofer*, *M. guilliermondii*, *S. paludigena* and *Y. lipolytica*. We conclude that PRs can be used as a raw material for microbial growth, ethanol and SCO production, which is of economic and environmental importance.

## 1. Introduction

Agriculture is a very important economic activity that boosts prosperity and contributes to tackling poverty in many countries. According to The World Bank collection, agricultural activity worldwide in 2017 accounted for 3.43% of the gross domestic product (GDP). Specifically for Greece, in 2018 the agricultural domestic product represented 3.72%, of GDP, twice as much as in other European countries, while together with processing, nowadays exceeds 8% (World Bank, September 2020). However, the expansion and modernization of the agricultural sector, and especially the processing of agricultural products, has caused environmental concerns, due to the inevitable co-production of residues and wastewaters, which are produced in huge quantities annually. Besides, water use in agriculture accounts for 70% of total water use globally.

Two different types of agro-industrial residues exist, namely field plant residues, which include remaining plant parts (e.g. leaves, straw, stalks, seeds, roots etc.), and industrial residues, which include organic matter or plant parts produced by food and fruit industries (e.g. molasses, peels, fruit marc and seeds and fruit pulp) (Sadh et al., 2018). Among residues, lignocellulosic biomass is the most abundant residue worldwide, and although it can be considered as a promising cost-effective fermentation feedstock (Diwan and Gupta, 2019; Valdés et al., 2020b, 2020a), is treated as waste in many countries (Arevalo-Gallegos et al., 2017). The uncontrollable disposal of agro-industrial residues, especially those produced during industrial processing, to the landfills without any treatment, can cause environmental pollution on the surrounded areas and harmful effects on human and animal health (Ghinea et al., 2019).

The use of the lignocellulosic material as a substrate in biotechnological applications requires additional pretreatment phases, to make it fermentable (Caporusso et al., 2021; Mosier, 2005; Sarris and Papanikolaou, 2016; Valdés et al., 2020a), which increases the cost of the whole process. Lignocellulosic biomass consists of cellulose, hemicellulose and lignin. During the pretreatment procedure, lignin “seal” is removed and cellulose is released and becomes accessible for hydrolysis (Haghighi Mood et al., 2013; Sánchez and Cardona, 2008). Pretreatment methods are categorized in four groups, i.e. physical (i.e. grinding and milling, microwave irradiation, mechanical extrusion, freeze pretreatment), chemical (in an acid or an alkaline environment), physico-chemical (using steam explosion, hot water, wet oxidation and CO_2_ explosion) and biochemical (using microorganisms and enzymes) (Haghighi Mood et al., 2013; Sarkar et al., 2012). Lately, combination of two of the above mentioned methods is proposed to increase the efficiency of the process. The first step of pretreatment involves lignin destruction, followed by hydrolysis of cellulose using acids or enzymes, resulting in the release of fermentable sugars (Sánchez and Cardona, 2008). Usually, glucose and xylose are extracted from lignocellulosic biomass. Thus, microorganisms that can utilize both C-5 and C-6 sugars are highly desired to increase the fermentation efficiency for the production of valuable metabolic products, including ethanol and lipids, from such materials (Diwan and Gupta, 2020).

The last decade the popularity of pomegranate products (mainly fresh fruit and juice) has increased tremendously thanks to their nutritional value and benefits to human health. Many reports describe the antimicrobial, antiviral, anticancer, antioxidant and antimutagenic properties of the various parts of the plant. (Akhtar et al., 2015; Bassiri-Jahromi, 2018; Hasnaoui et al., 2014; Joseph et al., 2013; Sharma et al., 2017). As a result, pomegranate cultivation has increased significantly worldwide to meet the growing demand. For instance, in Greece this crop extended up to 700-800 acres until 2007 and nowadays has reached 15.000 acres, yielding 30.000 tons of fruits per year. Consequently, the pomegranate processing industry has significantly expanded, and its negative outputs have already appeared (Kaderides et al., 2015). It has been estimated that the production of each ton of concentrated to 65 Brix pomegranate juice generates up to 5–5.5 tons of pomegranate residues (PRs) (i.e. pomegranate peels and seeds) (Hasnaoui et al., 2014). Although PRs is an excellent source of sugars, such as ribose, glucuronic acid, galacturonic acid, L-rhamnose, D-fucose, L-arabinose, D-xylose, D-mannose, D-galactose and D-glucose (Zhai et al., 2018a, 2018b) and numerous bioactive compounds (Santos et al., 2019; Sood and Gupta, 2015), in practice they are usually left unexploited and only a small amount is used as a soil conditioner (after composting), as fuel (in the form of dry pellets), as animal feed or for the recovery of pharmaceuticals and cosmetics (Goula et al., 2017; Pathak et al., 2017; Pereira et al., 2016; Pocan et al., 2018; Siddiqui et al., 2019). On the other hand, the necessity for low-cost and environmentally friendly substrates suitable for microbial processes related to the production of high value-added products (i.e. biofuels, single cell oils, single cell protein, enzymes, pigments, organic acids, etc.), has been arisen.

Biofuels (mainly bioethanol and biodiesel) are renewable, cost-effective and environmentally friendly energy sources alternative to fossil fuels, which has attracted the interest of many countries in the context of the diminishing oil dependence and eliminating the adverse effects of its use on global warming. Therefore, the demand for biofuels is gradually increasing worldwide. Specifically, bioethanol apart from its variety of commercial applications in cosmetology and in food and pharmaceutical industry, is also widely used as an engine fuel, on its own, or blended with gasoline. However, the utilization of the so-called 1_st_ generation” bioethanol, deriving from the fermentation of hydrolyzed corn starch or sucrose juice, runs into the “food *vs* fuel” dilemma (Gray et al., 2006), therefore the production of the so-called “2^nd^ generation” bioethanol (deriving through the valorization of waste streams) is considered as a very important scientific priority worldwide (Rastogi and Shrivastava, 2017; Sarkar et al., 2012; Tian and Chen, 2016). On the other hand, biodiesel produced by a variety of renewable resources (i.e. plant or animal fats) (Banković-Ilić et al., 2014; Gutiérrez et al., 2017), is non-toxic, biodegradable, and has a favorable emission profile, but, being potentially competitive in food production, it faces similar criticism to that of the “1_st_ generation” bioethanol. Alternatively, waste fats can be considered, with some restrictions, as a raw material for the production of biodiesel, and this option has aroused great interest in many countries. Besides, microbial oil, so-called single cell oil (SCO), produced from yeasts cultivated on agro-industrial residues, presenting a fatty acid (FA) composition similar to that of common vegetable oils, could be considered shortly as feedstock in the biodiesel manufacture (Dourou et al., 2018; Li et al., 2008; Papanikolaou and SCOs, fungi- and microalgae-derived SCOs are of pharmaceutical and dietary interest thanks to the presence of polyunsaturated fatty acids (PUFAs), such as γ-linolenic acid (GLA, C18:3, n-6), stearidonic (SDA, C18:4, n-3), arachidonic acid (ARA, C20:4, n-6), eicosapentaenoic acid (EPA, C20:5, n-3), docosapentaenoic acid (DPA, C22:5, n-3) and docosahexaenoic acid (DHA, C22:6, n-3) (Bellou et al., 2016; Fakas et al., 2009; Slaný et al., 2021).

A significant part of the production costs of both ethanol and SCO refers to the substrate cost suggesting that a choice of a low-cost material as substrate, such as a carbon-rich agro-industrial residue, is crucial for the process sustainability (Dourou et al., 2016; Osorio-González et al., 2019; Sarris et al., 2013). Moreover, new strategies using microorganisms derived after genetic engineering and/or adaptive laboratory evolution approaches, used to increase the efficiency of substrate assimilation and metabolite production, can contribute to the success of the process (da Silveira et al., 2020; Daskalaki et al., 2019; De Melo et al., 2020; Dourou et al., 2018; Li et al., 2019).

The aim of the current investigation was to study the conversion of PRs into high added value microbial products, i.e. ethanol and SCOs, using technologically suitable microbial strains in an integrated approach, which includes an efficient and environmentally friendly pretreatment followed by the production of microbial metabolic products, while minimizing management issues caused by PR disposal into the environment. The yeasts *Candida tropicalis*, *Lipomyces lipofer*, *Meyerozyma guilliermondii*, *Saccharomyces cerevisiae*, *Scheffersomyces coipomoensis*, *Sugiyamaella paludigena* and *Yarrowia lipolytica* were tested for their ability to grow on PR extract and produce ethanol and/or SCOs. Solid state fermentations (SSFs) of the fungus *Cunninghamella echinulata* were performed using PRs alone or in combination with wheat bran (WB) or oat flakes (OFs) as substrate. The final fermented product was enriched with polyunsaturated FAs, thus having a potential value used in animal or human nutrition. We conclude that PRs and PR extract are suitable microbial substrates and can be used as raw material in various biotechnological applications.

## 2. Material and Methods

### 2.1 Microorganisms and culture conditions

The following yeast strains were used: *Saccharomyces cerevisiae* AXAZ-1, *Candida tropicalis* NRRL Y-12968, *Lipomyces lipofer* NRRL Y-11555, *Yarrowia lipolytica* ACA DC 50109 (culture collection of Agricultural University of Athens, Greece) and three newly isolated strains from Chilean Valdivian Forest namely *Meyerozyma guilliermondii* ACA-DC 5397, *Scheffersomyces coipomoensis* ACA-DC 5395, *Sugiyamaella paludigena* ACA DC 5396 (Valdés et al., 2020b). In addition, the fungal strain *Cunninghamella echinulata* CCF 2195 (culture collection of fungi, Charles University, Prague, Czech Republic) was used in this research. All microbial strains were maintained at −80 °C in glycerol 30% and for short term maintenance on potato dextrose agar (PDA, Himedia, Mumbai, India) slants at 4 ± 1 °C and regularly sub-cultured.

### 2.2 PR collection, media preparation and culture conditions

Fresh PRs (i.e. pomegranate peel and seeds remaining after juice extraction, cultivar *Wonderful*) were obtained from the Bacfresh Company located in Achaia regional unit of Greece, or produced in the laboratory after juice extraction using a mechanical home juicer. PR moisture was gravimetrically determined after drying at 105 °C in an oven until constant weight. Polysaccharide extraction and hydrolysis were simultaneously performed from wet PRs cut into pieces of 0,5 × 0.5 cm in water or in H_2_SO_4_ (Fluka, Steinheim, Germany) solution used at a ratio of 40 mL of water or H_2_SO_4_ solution to 10 g of PRs. The treatment was performed using various concentrations of H_2_SO_4_ (up to 0.10 M), at temperature 100 or 120 °C, pressure 101 or 219 kPa and time ranging from 20 to 120 min. The treated PR mixture was subsequently centrifuged (Heraeus, Biofuge Stratus, Thermo Scientific, Osterode, Germany) at 19,000 × g, 4 °C for 10 min and the supernatant, optionally diluted with water, was used as culture medium. Supernatant pH was determined using an ORION pH meter (Boston, USA) and adjusted before inoculation to the desirable level (ranged from 2.6 to 6.5 ± 0.1) by adding 4 M NaOH (Merck) or 4 M HCL (Sigma-Aldrich).

Submerged cultures under near anaerobic conditions were conducted by yeast strains in manually periodically agitated 200 mL Duran flasks (Bioengineering, Wald, Switzerland) containing 50 ± 1 mL of PR extract. The flasks were incubated in a Memmert GmbH+Co. KG, Germany incubator at temperature 28 °C. Submerged cultures under aerobic conditions were performed in 250 mL Erlenmeyer flasks containing 50 ± 1 mL of PR extract incubated in a rotary shaker (ZHICHENG ZHWY 211C, Shanghai, China) at agitation rate 180 rpm and temperature 28 °C.

After medium sterilization (at 121 °C for 20 min) the flasks were inoculated with 1 mL of a pre-culture (containing 10^8^ cells), carried out in a medium containing (in g/L): yeast extract (Conda) 3; glucose (AppliChem, Darmstadt, Germany) 10; peptone (Himedia) 5; malt extract 3 (Sigma-Aldrich, Steinheim, Germany), and incubated in a rotary shaker as above.

SSFs of wet chopped peels alone or in mixture with wheat bran (WB) or oat flakes (OFs) [used in different proportions to control moisture (Klempová et al., 2020)] were performed by *C. echinulata* in high-density polyethylene (HDPE) bags (20 × 30 cm) containing 10 g of substrate at room temperature (T = 25 ± 2 °C). HDPE bags ensure aseptic conditions and smooth oxygen transfer during fermentation. After sterilization (at 105 °C for 45 min) the bags were inoculated with 10^7^ spores/mL obtained from a 7-day-old culture grown on polished rice (Slaný et al., 2020). After inoculation the bags were closed and the substrate was arranged on the shelves to obtain a substrate layer thickness of around 5c:mm. SSFs were carried out under static conditions for up to 5 days.

### 2.3 Microbial growth and biomass determination

Yeast growth was estimated by enumerating the number of cells/mL using a haemocytometer cell counting chamber (Neubauer improved, Poly-Oprik, Bad Blankenburg, Germany). Dry cell mass (x, g/L) was gravimetrically determined after harvesting of yeast cells by centrifugation at 19,000 × g for 15 min at 4 °C. The cells were washed twice with NaCl 0.9% (w/v) and dried at 80 °C until constant weight.

The fermented substrate after SSFs (around 10 g) was washed with 60 mL of distilled water and filtrated through Whatman No. 1 paper. The obtained byproduct was gravimetrically determined after drying at 60 °C until constant weight. The concentration of reducing sugars (R.S.) was determined in the filtrate (see below) and the fungal growth was indirectly estimated by calculating R.S. consumption. Besides, GLA concentration in lipids (estimated as described below) of the fermented substrate was additionally used as a marker to indirectly estimate fungal growth, as this FA is absent from the substrate.

### 2.4 Sugar and phenolics determination

The concentration of total sugars (T.S.) and reducing sugars (R.S.) in the liquid growth medium and in the solid substrate extract was determined according to Dubois et al. (1951) and DNS (Miller, 1959) methods, respectively, and expressed as glucose equivalent (in g/L). Sugar concentrations were also expressed per weight of dry substrate. The total phenolic compounds in the liquid growth medium and in the solid substrate extract were determined according to Folin and Ciocalteau (1927). The concentration of phenolic compounds was expressed as gallic acid equivalent (in mg/L or per weight of dry substrate).

### 2.5 Ethanol determination

Ethanol was determined in filtered aliquots of the culture supernatant by high performance liquid chromatographer (HPLC - Ultimate 3000 Dionex, Germering, Germany) equipped with a Reflective Index detector (RI-101, Shodex, Kawasaki, Kanagawa Japan) and an AminexHPX-87H column (300 × 7.8 mm). H_2_SO_4_ (Fluka) 0.005 N was used as eluent at a flow rate of 0.6 mL/min. The column temperature was 65 °C.

### 2.6 Lipid extraction and purification

Lipids from dry yeast cell mass or dry homogenized fermented substrate were extracted in chloroform:methanol (Sigma-Aldrich) (2:1 v/v) according to Folch et al. (1957) method as modified by Dourou et al. (2017). The extracts were filtered through Whatman No. 1 paper, washed with a KCl (Sigma-Aldrich) 0.88% (w/v) solution and dried over anhydrous Na_2_SO_4_ (Sigma-Aldrich). Finally, the solvents were evaporated under vacuum using a Rotavapor R-20 device (BUCHI, Flawil, Switzerland), and the total lipids were gravimetrically determined.

### 2.7 Fatty acid composition of microbial lipids

FA moieties of total lipids were converted into their FA methyl esters (FAMEs) in a two-stage reaction according to AFNOR. FAMEs were analyzed in a GC apparatus (Agilent 7890 A, Agilent Technologies, Shanghai, China), equipped with a flame ionization detector (FID) and an HP-88 (J&W Scientific) column (60 m × 0.32 mm). Helium was used as carrier gas at a flow rate of 1 mL/min. Injection temperature was 250 °C, the oven temperature was 200 °C and FID temperature was 280 °C. Peaks of FAMEs were identified by reference to authentic standards.

### 2.8 Thin-layer chromatography (TLC) analysis

Known weight of total lipids (approximately 300 mg) were loaded in the form of narrow bands of 1 cm on Merck TLC silica gel 60 plates using a CAMAG ATS 4 apparatus and separated by a developing system for non-polar lipids using a solvent system consisting of hexane/diethyl ether/acetic acid (80:20:1, v/v/v). TLC plates were visualized using a combustion water solution containing 3.3% (v/v) sulfuric acid, 50% (v/v) methanol, 0.33% (w/v) MnCl_2_.4H_2_O, and dried at 130 °C for 10 min. After visualization, TLC plates were scanned by CAMAG TLC Scanner 4 at 400 nm and evaluated by winCATS software ver. 1.4.8 (CAMAG) to evaluate lipidic structures such as TAGs, DAGs, FFAs, etc. in lipid extract (Gajdoš et al., 2015).

### 2.9 Modelling

The Hinshelwood model (Hinshelwood, 1947) was used to describe yeast growth and ethanol production under aerobic and anaerobic conditions. The balance equations for cell mass and ethanol production and sugar consumption are:

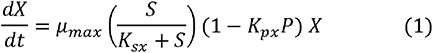

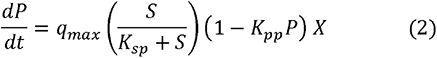

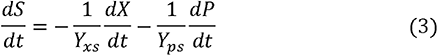

where, *X*, *S* and *P* are cell mass, sugars and ethanol concentrations (g/L), respectively, is the fermentation time (h), *μ*_*max*_ is the maximum specific growth rate (1/h), *q*_*max*_ is the maximum specific ethanol production rate (1/h), *K*_*sx*_ and *K*_*sp*_ are the saturation constants (g/L), *K*_*px*_ and *K*_*pp*_ are the ethanol inhibition constants (g/L), *Y*_*xS*_ is the yield coefficient for biomass production with respect to sugars (g biomass/g sugars) and *Y*_*pS*_ is the yield coefficient for ethanol production with respect to sugars (g ethanol/g sugars).

The fitting of the model equations on the experimental data, was performed using the numerical code Aquasim (Version 2.1d), which uses a fully implicit finite difference spatial discretization Gear scheme in conjunction with the algorithm DASSL. Aquasim uses the weighted least-squares method to estimate the optimum parameter values (Reichert, 1998).

### 2.10 Statistical analysis

The experimental data were treated using OriginPro 2018 Software (OriginLab Corporation, Northampton, MA, United States). One-way analysis of variance ANOVA followed by a Bonferroni post hoc test was performed to determine significant differences. The null hypothesis was rejected at a significance level of p ≤ 0.01 or 0.05.

## 3. Results and Discussion

### 3.1 Pomegranate juice extraction and composition of the PRs

The composition of the PRs depends not only on the cultivation conditions and the cultivar but also on the efficiency of the extraction of the juice and the extraction technology used (Catania et al., 2020; Mphahlele et al., 2016). Thus, compositional differences concerning mostly initial sugar and phenolic compounds concentration were observed among samples, which were used in the current investigation. The PRs resulting from the processing of the fruit under laboratory conditions constitute 68.7 ± 1.9% of the total weight of the fruit, while this percentage can reach up to 80% under industrial processing conditions. Consequently, the yield in juice under laboratory conditions was 29.3 ± 1.3%, while under industrial conditions it is estimated to be less than 20%. The presence of phenolics and flavonoids in the pomegranate juice is indeed suitable, due to the beneficial effects of these compounds on human health. However, exhaustive fruit juicing is not generally selected by the industry to avoid the exceeding enrichment of the juice in such compounds, which may affect the acidity and the color of pomegranate juice and, therefore, its quality (Catania et al., 2020; Türkyılmaz et al., 2013). The moisture of the solid residue produced under laboratory and industrial conditions was 66% and 75%, respectively.

### 3.2. Pretreatment of the PRs

PRs were cut into small pieces to reduce their size, increase the accessible area and improve the porosity. Following, PRs were treated for 20-120 min under acidic conditions with a H_2_SO_4_ solution up to 0.10 M, at high temperature (i.e. 100 and 120 °C) and pressure (i.e. 101 and 213 kPa) for the simultaneous extraction and hydrolysis of the polysaccharides (Table 1). The efficiency of the extraction process was evaluated by calculating the percentage of T.S. extracted in the solution on the T.S. extracted under the strongest conditions (i.e. temperature 121 °C, pressure 219 kPa, H_2_SO_4_ 0.10 M, for 60 min). Moreover, the efficiency of the hydrolysis process in each case was evaluated by calculating the percentage of R.S. on the T.S. in the extract.

**Table 1:** Polysaccharide extraction and hydrolysis from PRs under various experimental conditions of H_2_SO_4_ concentration, processing time and temperature. The asterisk (*) indicates one set experiments performed using an open vessel assuring the presence of O_2_. Data are presented as mean values of three replications. Letters refer to comparisons at vertical reading only; different letters indicate that differences of means are statistically significant at p ≤ 0.05. Data were treated by one-way ANOVA test followed by Bonferroni post hoc test.

| Set | Time (min) | Temperature (°C) | H<sub>2</sub>SO<sub>4</sub> (M) | Total Sugars (g/L) | Reducing Sugars (g/L) | Extraction (%) | Hydrolysis (%) |
| --- | --- | --- | --- | --- | --- | --- | --- |
| a | 20 | 121 | 0.00 | 34.7<sup>a</sup> ± 2.3 | 28.3<sup>a</sup> ± 0.9 | 100.6 | 81.6 |
| b | 30 | 121 | 0.00 | 30.3<sup>a</sup> ± 1.6 | 29.0<sup>a</sup> ± 1.6 | 86.6 | 95.7 |
| c | 60 | 121 | 0.00 | 32.5<sup>a</sup> ± 2.6 | 29.8<sup>a</sup> ± 2.6 | 94.2 | 91.7 |
| d | 60 | 121 | 0.03 | 32.5<sup>a</sup> ± 2.2 | 31.6<sup>a</sup> ± 3.0 | 94.2 | 97.2 |
| e | 60 | 121 | 0.05 | 33.5<sup>a</sup> ± 2.7 | 32.5<sup>b</sup> ± 3.4 | 97.1 | 97.0 |
| f | 60 | 121 | 0.07 | 30.4<sup>a</sup> ± 2.0 | 28.5<sup>a</sup> ± 3.3 | 88.1 | 93.8 |
| g | 60 | 121 | 0.10 | 34.5<sup>a</sup> ± 1.9 | 28.6<sup>a</sup> ± 3.1 | 100.0 | 82.9 |
| h | 120 | 121 | 0.00 | 30.9<sup>a</sup> ± 0.8 | 31.1<sup>a</sup> ± 0.3 | 89.6 | 100.6 |
| i | 120 | 100 | 0.00 | 31.7<sup>a</sup> ± 0.2 | 26.1<sup>a</sup> ± 0.6 | 91.9 | 82.3 |
| j* | 120 | 100 | 0.00 | 30.1<sup>a</sup> ± 0.5 | 24.9<sup>b</sup> ± 0.2 | 87.2 | 82.7 |

Acid pretreatment is recommended for hardwoods and agricultural residues, while the kind of acid used (e.g. HCl, H_2_SO_4_, HNO_3_ etc.), its concentration, temperature and reaction time differ in the various protocols described in the literature (Nitsos et al., 2018; Zhang and Bao, 2018). In the current investigation, we have demonstrated that the addition of a strong acid to PRs is not necessary to achieve adequate extraction of polysaccharides and their hydrolysis to R.S., probably due to the high natural acidity of PRs imparting a pH ≤ the best physicochemical properties of water, as an extractant, against sulfuric acid solutions. Specifically, the extraction efficiency achieved in water at temperature 121 °C, under 219 kPa pressure, for 60 min was no statistically different (at p=0.01 or 0.05) to that achieved under similar conditions in the presence of H_2_SO_4_ used at concentrations 0.03-0.10 M (comparisons of entries c-g, Table 1). Similarly, the utilization of H_2_SO_4_ did not significantly increase the hydrolysis efficiency (comparisons of entries c-g, Table 1). It is reported that the extraction efficiency of ingredients increases with fluidity and, in some cases, with solvent polarity (Xu et al., 2019). Low viscosity of the solvent is generally required to obtain high extraction yields since high viscosity hinders the mass transfer of the solvent to the target solutes. High polarity solvents are also required when polar molecules, such as polysaccharides, are targeted for extraction. Therefore, the higher polarity of water in combination with its low viscosity could explain the higher extraction yields obtained in this research in the absence of sulfuric acid. Çam and Hişil (2010) investigating polyphenols extraction from pomegranate peels, reported that water extraction was as effective as conventional methanol extraction.

The processing time is another factor that influences the extraction efficiency (Zhu and Liu, 2013). Comparing the effect of the processing time on polysaccharide extraction under the above-mentioned conditions (i.e. in water, at T=121 °C, P=219 kPa), we ascertain that the extraction yield at 20 min was higher compared to higher processing time (i.e. 30, 60, and 120 min, entries a-c and h in Table 1), although not statistically significant at p=0.01 and 0.05. This apparent reduction in extraction yield is probably a result of sugar degradation. Therefore, the extraction time of 20 min was selected as a sufficient time for the extraction and hydrolysis of polysaccharides from PRs.

Trying to reduce the processing temperature more experiments were performed at 100 °C and atmospheric pressure (i.e. 101 kPa) instead of 121 °C and 219 kPa. We note that the extraction efficiency was higher and the efficiency of hydrolysis lower at 100 °C than at 121 °C (comparisons of entries h and i, Table 1). Zhu and Liu (2013) examining the effect of different temperatures (from 70 to 100 °C) on polysaccharide extraction, reported that maximum yield was obtained at 95 - 100 °C. Finally, at T=100 °C comparing the effect of O_2_ presence (entries i and j in Table 1), both T.S. and R.S. yields (in g/L) were higher (i.e. 5.3 and 4.8% respectively) when a closed system, almost in the absence of O_2_ was applied.

Generally, high temperature increases solubility and diffusion of the ingredients to be extracted and decreases the viscosity of the solvent, which is intended. Though, pressure seems to have a great influence on the extraction and hydrolysis yield too. Comparing the entries h-j in Table 1, it seems that conditions reported in entry i would be appropriate. However, the high processing time is not desirable, therefore, the conditions that we conclude to use for this research are those reported in entry a: in water, for 20 min, at T=121 °C and P=219 kPa.

The selection of the pretreatment method of lignocellulosic biomass and its operating conditions are fundamental for a successful polysaccharide extraction and hydrolysis, and the subsequent fermentation. Methods of chemical pretreatment such as acid, alkaline and their combination, are the most preferred techniques in recent years, thanks to their flexibility of applications at the industrial level, and the high final yields of assimilable sugars achieved (Valdés et al., 2020a). Among them, acid pretreatment, and specifically diluted acid pretreatment, is favorable. However, regardless of the pretreatment method used, toxic compounds are generated as by-products, such as formic, acetic and levulinic acid, neutral and acidic phenolics, and other chemicals in hydrolysates, which can inhibit microbial growth (Bravo et al., 2017; Haghighi Mood et al., 2013; Janga et al., 2012; Yu et al., 2018).

### 3.3 Microbial growth on PR based media

The growth of microorganisms on PRs presents various peculiarities related to the nature of the substrate, including assimilability of the contained carbon sources, the presence of microbial inhibitors (i.e. natural such as phenolic compounds or generated during pre-treatment), and its low pH values. Therefore, the challenge is to identify microbial strains capable of growing on PRs and PR extract. In the present study, the ability of the oleaginous fungus *C. echinulata* and the yeasts *C. tropicalis*, *L. lipofer, M. guilliermondii, S. cerevisiae*, *S. coipomoensis*, *S. paludigena* and *Y. lipolytica* to grow on PRs or PR extract and produce various metabolites was studied under several growth conditions (Table 2). To the best of our knowledge, the current investigation is the first report in which both pomegranate peels and seeds have been employed as fermentation medium for microbial growth, while only recently, pomegranate peels have been treated as a substrate for biodiesel production by *Bacillus cereus* (Kanakdande et al., 2020) and for citric acid production by the fungus *Aspergillus niger* (Roukas and Kotzekidou, 2020).

**Table 2:** Cell mass, ethanol production, lipid accumulation and phenolic removal by different microorganisms cultivated on PR extract or solid residue under different culture conditions. Abbreviations: R.S., reducing sugars; nd, not determined; ND, not detected

| Microbial strain | Culture conditions |  |  | Initial R.S.<br>concentration<br>(g/L or mg/g<br>by-product) | X<sub>max</sub><br>(g/L) | Ethanol<sub>max</sub><br>(g/L) | L/X<sub>max</sub><br>% | Initial phenolic<br>concentration<br>(g/L or mg/g by-product) | Phenolic<br>removal<br>(%, w/w) |
| --- | --- | --- | --- | --- | --- | --- | --- | --- | --- |
|  | Fermentation | Growth | pH |  |  |  |  |  |  |
| *C. echinulata* | Solid state | Aerobic | 6.5 | 163.7<sup>a</sup> | nd | - | 11.7 | 14.0 | 16.8 |
|  |  |  | 6.5 | 22.3<sup>b</sup> | nd | - | 4.8 | 0.8 | 44.3 |
|  |  |  | 6.5 | 7.5<sup>c</sup> | nd | - | 7.1 | 2.5 | 59.9 |
| *C. tropicalis* | Submerged | Aerobic | 4.5 | 37.0<sup>d, #</sup> | nd | 4.9 | - | nd | nd |
|  |  | Anaerobic | 4.5 | 32.1<sup>d, *</sup> | nd | 6.8 | - | nd | nd |
| *L. lipofer* |  | Aerobic | 6.5 | 22.5<sup>d, #</sup> | 2.1 | - | 7.0 | 1.6 | 29.8 |
|  |  |  | 6.5 | 15.3<sup>e, #</sup> | 1.6 | - | 3.9 | 0.9 | 18.3 |
|  |  |  | 6.5 | 9.6<sup>f, #</sup> | 1.0 | - | 13.5 | 0.5 | 36.5 |
| *M. guilliermondii* |  | Aerobic | 2.6 | 18.0<sup>d, #</sup> | 2.1 | 3.2 | 18.1 | 2.3 | 48.8 |
|  |  | Anaerobic | 2.6 | 19.3<sup>d, #</sup> | 1.7 | 5.8 | 4.7 | 2.3 | 22.3 |
| *S. cerevisiae* |  | Anaerobic | 3.5 | 30.8<sup>d, #</sup> | 2.7 | 7.4 | - | 2.6 | ND |
|  |  |  | 3.5 | 37.4<sup>d, *</sup> | 2.8 | 12.5 | - | nd | nd |
|  |  |  | 3.5 | 32.3<sup>d, *</sup> | 2.7 | 9.3 | - | nd | nd |
|  |  |  | 5.0 | 33.3<sup>d, #</sup> | 2.8 | 9.7 | - | 2.5 | ND |
| *S. coipomoensis* |  | Aerobic | 2.6 | 20.8<sup>d, #</sup> | 0.7 | 0.2 | 8.3 | 2.0 | 22.7 |
|  |  | Anaerobic | 2.6 | 21.5<sup>d, #</sup> | 1.0 | 3.6 | 4.2 | 2.0 | 30.6 |
| *S. paludigena* |  | Aerobic | 2.6 | 18.0<sup>d, #</sup> | 2.1 | 1.6 | 15.4 | 2.3 | 61.0 |
|  |  | Anaerobic | 2.6 | 19.3<sup>d, #</sup> | 1.5 | 5.3 | 9.0 | 2.3 | 22.1 |
| *Y. lipolytica* |  | Aerobic | 6.5 | 21.8<sup>d, #</sup> | 1.7 | - | 7.0 | 2.6 | 45.0 |
|  |  |  | 6.5 | 7.6<sup>f, #</sup> | 0.9 | - | 11.0 | 1.3 | 64.5 |

<sup>a</sup>Fermentation on 100% PRs; <sup>b</sup>Fermentation on blend 10% PRs and 90% WB; <sup>c</sup>Fermentation on blend 20% PRs and 80% OFs; <sup>d</sup>100% PR extract; <sup>e</sup>80% PR extract and 20% water; <sup>f</sup>50% PR extract and 50% water; <sup>#</sup>laboratory processing conditions; <sup>\*</sup> industrial processing conditions

A sustainable process for microbial conversion of lignocellulosic biomass into biofuels and other products is based on the capability of the microbial strains used to assimilate/co-assimilate, the various sugars (including C-5 and C-6) released during pretreatment (Valdés et al., 2020a). All microbial strains tested in the current investigation were able to grow on PRs or PR extract showing different biochemical potential and effectiveness (Table 2). Among them, the new isolates from the Chilean Valdivian forest identified as *M. guilliermondii*, *S. coipomensis*, and *S. paludigena* recently characterized for their ability to grow on lignocellulosic hydrolysate-model media (Valdés et al., 2020b), showed interesting ethanol production and lipid accumulation capacity, cultivated under anaerobic and aerobic conditions, respectively. The most commonly used yeast for ethanol production is *S. cerevisiae*. However this yeast is not able to metabolize pentose sugars naturally, such as xylose and arabinose, to ethanol, and therefore several strategies on engineering pentose metabolism on *S. cerevisiae* have been proposed (Fernandes and Murray, 2010; Gopinarayanan and Nair, 2019; Hahn-Hägerdal et al., 2007). Interestingly, *S. cerevisiae* during submerged cultures on PR extract in this study showed remarkable growth and ethanol production ability. Contrary, growth and ethanol production by *C. tropicalis* was not satisfactory.

*L. lipofer* and *Y. lipolytica* were both able to accumulate lipids during submerged cultures on PRs extract. Strains belonging to the genus of *Lipomyces* have notable oleaginous capacity and are suitable for SCO production from lignocellulosic biomass and other agro-industrial wastes (Di Fidio et al., 2020; Dien et al., 2016; Dourou et al., 2016; Gong et al., 2012; Liang and Jiang, 2013; Sitepu et al., 2013; Slininger et al., 2016; Vasaki et al., 2021; Zhao et al., 2008). *Y. lipolytica*, the most studied oleaginous yeast, is able to grow on a variety of agro-industrial substrates and to produce high added value metabolites (Carota et al., 2020; Chatzifragkou et al., 2011; da S. Pereira et al., 2019; Dobrowolski et al., 2016; Dourou et al., 2016; Makri et al., 2010; Patsios et al., 2020). Over the last decade, a variety of studies have focused on modifications of *Y. lipolytica* genome through genetic engineering to enable the efficient assimilation of sugars found in lignocellulosic biomass (Ledesma-Amaro and Nicaud, 2016; Niehus et al., 2018; Yook et al., 2020).

The oleaginous fungus *C. echinulata* was able to synthesize lipids during SSF on sole PRs or blends of PRs with WB or OFs (Table 2). WB and OFs are also produced in abundance, consisting yet again lignocellulosic residues which need management. In 2010, the National Green Tribunal established control measures to stop such biomass burning, due to air pollution, encouraging the exploitation of agricultural residues as a substrate for microbial production of high added-value products using environmentally friendly approaches. Filamentous fungi belonging to the *Cunninghamella* genus and related genera play an important role in developing sustainable biorefinery processes thanks to their ability to utilize a broad range of renewable feedstock, and waste materials (e.g. corn gluten, corn steep, orange peel and tomato waste hydrolysate) and convert them into SCOs containing significant quantities of GLA (Diwan and Gupta, 2019; Donot et al., 2014; Fakas et al., 2008; Gema et al., 2002).

Phenolic compounds present in agro-industrial residues are principally responsible for their phytotoxicity and microbial growth inhibition, while their breakdown is considered to be the limiting step during biotreatment (Aggelis et al., 2003; Tsioulpas et al., 2002). Moreover, phenolics can be released from lignin as by-products during the pretreatment of lignocellulosic biomass, depending on the parameters of the process, such as temperature and duration of the treatment. Phenolic compounds, when present in the growth media, penetrate biological membranes and cause loss of their integrity affecting cell growth and the whole fermentation process (Baral and Shah, 2014). Therefore, their degradation and/or removal prior to the fermentation process is of high biotechnological interest. In this investigation, the PR extract used as a growth medium for submerged cultures of *L. lipofer* and *Y. lipolytica* was diluted in some cases with water to reduce the initial phenolic concentration and the results were compared to those obtained on undiluted PR hydrolysate (Table 2, see letters d, e, f). In addition, phenolic compounds were reduced in the case of SSF of *C. echinulata* by the incorporation of cereals. Nevertheless, remarkable phenolic removal (up to 30%) was performed in some cases by *C. echinulata*, *L. lipofer*, *M. guilliermondii*, *S. paludigena* and *Y. lipolytica*. Benzene compounds are degraded by a few stains belonging to the phylum Ascomycota (e.g. the yeasts *Debaryomyces hansenii*, *L. starkeyi*, and *S. cerevisiae,* the fungus *Geotrichum candidum*, and the yeast-like *Aureobasidium pullulans*) and the phylum Basidiomycota (*Rhodotorula*, *Trichosporon cutaneum*) (Pasha and Rao, 2009).

Low pH media are generally suitable for large scale applications since they are self-protected against bacterial contamination. Indeed, in media having exceptionally low pH, sterilization can be omitted, and the maintenance of aseptic conditions is not necessary. Thus, the fermentation cost can be significantly reduced. In the present study, the strains *M. guilliermondii, S. coipomoensis*, and *S. paludigena* were able to grow at pH = 2.6, *S. cerevisiae* at pH = 3.5 and 5.0, *C. tropicalis* at pH = 4.5, while *C. echinulata*, *L. lipofer* and *Y. lipolytica* were cultivated at their optimal pH value of 6.5.

### 3.4 Bioethanol production

Bioethanol produced using low- or negative-acquisition cost substrates, such as carbon-rich agro-industrial residues, is at the forefront of biotechnological/industrial interest for many years. Bioethanol can be used as a raw material in the food, pharmaceuticals and cosmetics industry or as biofuel. During alcoholic fermentation, the sugars in the form of hexoses or pentoses are transformed into pyruvic acid (through the metabolic path of glycolysis), which under anaerobic conditions, is converted into ethanol and CO_2_. The strain AXAZ-1 of *S. cerevisiae*, as well as *M. guilliermondii*, *S. coipomoensis*, and *S. paludigena* were used in the current investigation and proved able to grow well in PR extract and to efficiently convert sugars to ethanol cultivated under anaerobic conditions (Fig. 1 and 2). The yeast *S. cerevisiae* is widely used worldwide in winemaking, in the brewery and baking industry, as well as in many other biotechnological applications, as it possesses key features, such us quick fermentation ability, genetic stability, tolerance to low pH environments, ethanol tolerance, osmotolerance and thermotolerance (Nevoigt, 2008; Sarris et al., 2014). PR extracts derived under laboratory and industrial processing conditions were used as a fermentation medium for *S. cerevisiae*. As mentioned above, the difference between these processes is that under industrial juicing the resulting extract had a higher sugar concentration. *M. guilliermondii*, is a promising species implicated in various biotechnological applications, including the conversion of pentoses to ethanol (Martini et al., 2016; Matos et al., 2014; Yan et al., 2021). Similarly, *Scheffersomyces* and *Sugiyamaella* genera are reported as xylose-fermenting genera containing species having an ethanol production capacity (Jia et al., 2020; Lopes et al., 2018; Morais et al., 2020; Sena et al., 2017). This is the first report that describes bioethanol production from *S. coipomoensis* and *S. paludigena* strains.

**Fig. 1:**
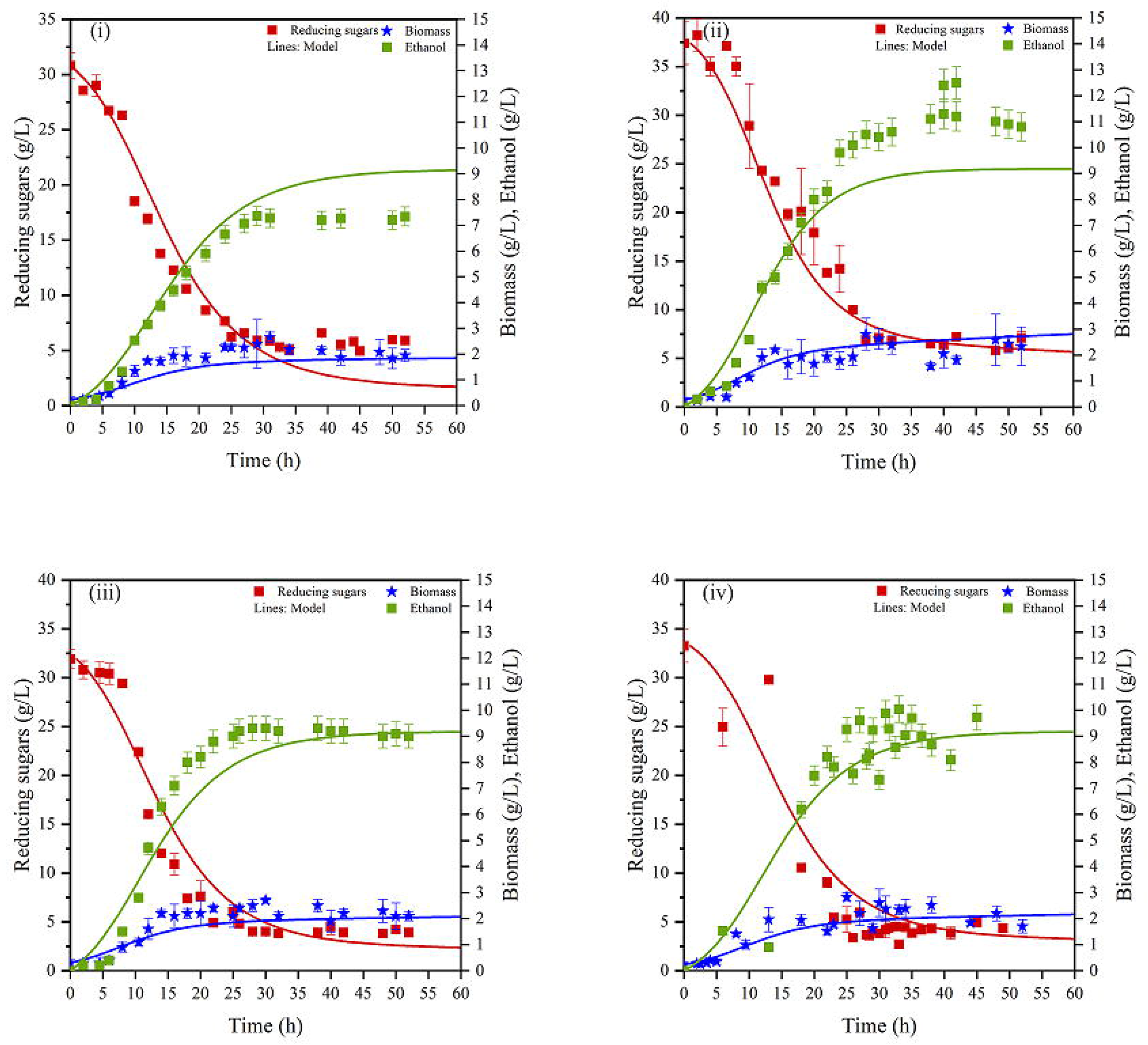
Kinetics of growth, ethanol production and substrate consumption of *Saccharomyces cerevisiae* under anaerobic conditions, cultivated at pH=3.5 and laboratory processing conditions (i), at pH=3.5 and industrial processing conditions (ii), (iii) and at pH=5.0 and laboratory processing conditions (iv). Points and lines represent experimental data and model prediction, respectively.

**Fig. 2:**
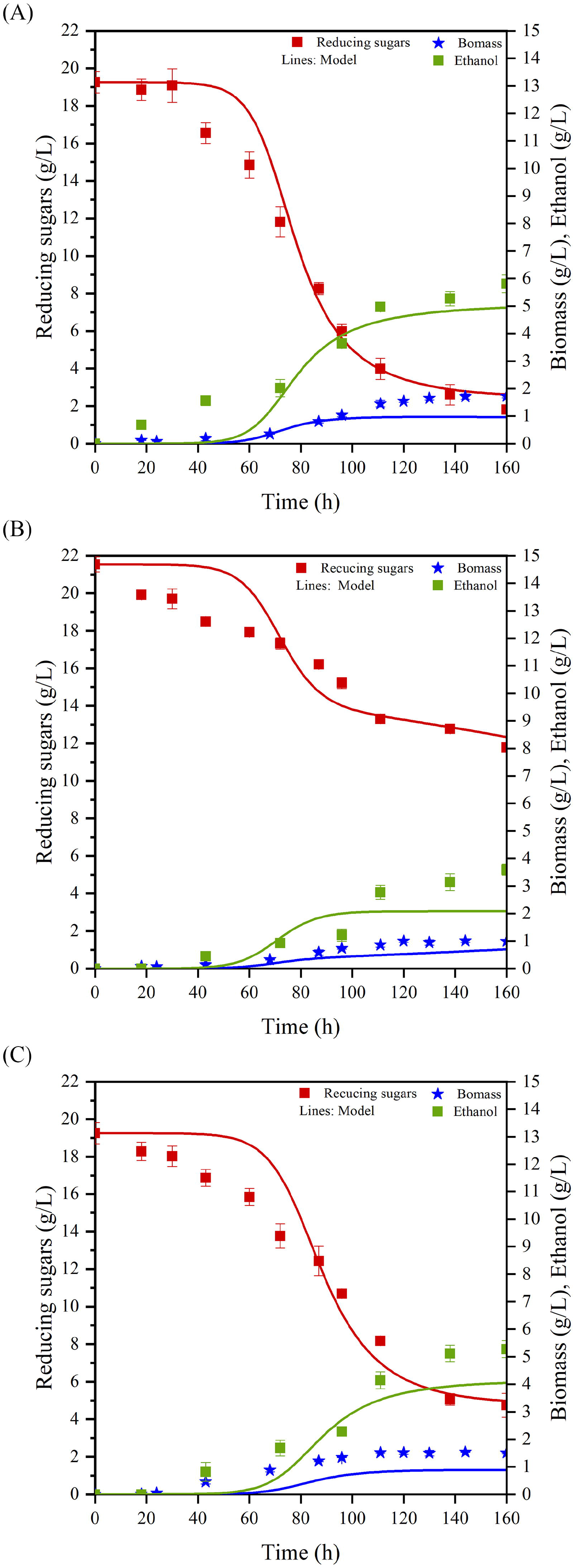
Kinetics of growth, ethanol production and substrate consumption of *Meyerozyma guilliermondii* (A), *Scheffersomyces coipomoensis* (B), and *Sugiyamaella paludigena* (C) cultivated under anaerobic conditions at pH=2.6 and laboratory processing conditions. Points and lines represent experimental data and model prediction, respectively.

Several mathematical models have been developed to describe the alcoholic fermentation process (Birol et al., 1998). Among them, the Monod (Monod, 1942) and Hinshelwood (Hinshelwood, 1947) models have been considered as suitable to describe the process under low/very low product inhibition (Birol et al., 1998; Kostov et al., 2012). In the current paper, the Monod model was initially employed to describe microbial growth, ethanol production and sugars consumption during fermentation and to calculate parameter values. However, it was observed that this model was not able to simulate the process for most of the yeast strains, probably due to the inhibitory effect of ethanol on growth and ethanol production (data not shown). Alternatively, the Hinshelwood model, which includes *K*_*px*_ and *K*_*pp*_ parameters indicating a week inhibition on cell growth and ethanol production by ethanol (Kostov et al., 2012), was employed to simulate the process. Fig. 1 and 2 show experimental data and model fitting, while the estimated values of the kinetic parameters along with the correlation coefficient (R^2^) are shown in Table 3.

**Table 3:**
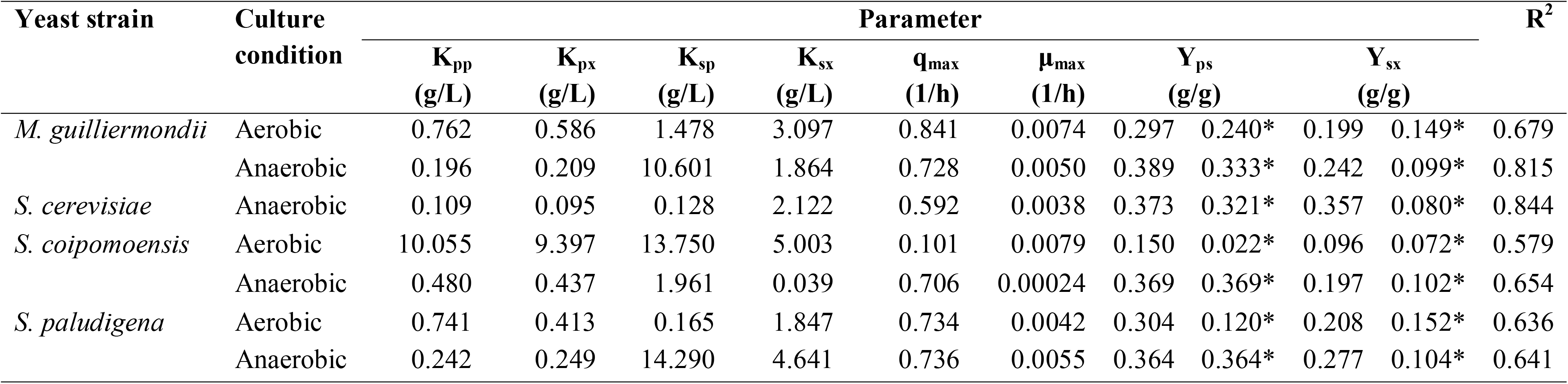
Kinetic parameter values of yeast strains growing under aerobic and anaerobic conditions in PRs extract. The asterisk (*) indicates the experimental values.

The *μ*_*max*_ values ranged between 0.0038 and 0.0079 1/h, except for the value obtained from the yeast *S. coipomoensis* in anaerobic conditions, which was 0.00024 1/h. The corresponding values in the literature using the Hinshelwood model range from 0.231 to 0.289 1/h by using free or immobilized cells of yeast *S. cerevisiae* growing on glucose (Kostov et al., 2012), sweet sorghum stalk juice (Jin et al., 2012) and corn stover (Tian and Chen, 2016). The low *μ*_*max*_ values in the present study may be due to the cell growth inhibition by ethanol and/or inhibitory compounds generated during PRs pretreatment. It has been reported that the presence in the fermentation medium of phenolics at 2.0 g/L negatively affects both cell viability of *S. cerevisiae* and ethanol production (Liu et al., 2016). In the present study, the initial concentration of phenolic compounds was about 2.0 g/L and remained almost constant during fermentation of *S. cerevisiae*. *Y*_*ps*_ calculated under anaerobic conditions ranged from 0.389 (for *M. guilliermondii*) to 0.364 (for *S. paludigena*), while the experimental values are close to those predicted by the model (Table 3). For *Y*_*sx*_, the values predicted by the model were in all cases higher compared to the experimental values for all yeast strains. These discrepancies may be due to the presence of growth inhibitors on PR extract that are not included in the Hinshelwood model. The estimated values of *K*_*pp*_, *K*_*px*_, *K*_*sx*_ and *K*_*sp*_ are in the range of values reported in the literature (Birol et al., 1998; Jin et al., 2012; Kostov et al., 2012; Tian and Chen, 2016).

In all experiments, the mathematical model adequately predicts the biomass and ethanol production as well as sugar consumption, which was confirmed by the high R^2^ values in most of the cases. The kinetic parameters values vary among strains indicating that each of them exhibits a different growth behavior on PR extract under aerobic and anaerobic conditions (Table 3). Most yeast strains produced ethanol ranged between 3.6 and 12.5 g/L showing high productivities evidenced by the high values of the parameter *q*_*max*_ (ranged h). Exceptionally, *S. coipomoensis* cultivated under aerobic c: conditions did not produce significant ethanol quantities (up to 0.186 g/L at the end of fermentation), thus low *q*_*max*_ value was observed. Kostov et al. (2012) found that *q*_*max*_, calculated using the Hinshelwood model fitted on the experimental data, of free cells of *S. cerevisiae* was 0.452 g/g h when cultivated on 110 g/L of glucose, while this value increased h using immobilized cells of *S. cerevisiae*.

Worthy of mentioning is that all fermentations were conducted under low pH values. Specifically, *S. cerevisiae* was cultivated at pH=3.5 (natural pH of PR extract) and pH=5.0 (after adjustment) on PR extract derived under laboratory processing conditions (Table 2), and ethanol and cell mass production was similar. Moreover, the pH during fermentation of *M. guilliermondii*, *S. paludigena* and *S. coipomoensis* was 2.6. The above-mentioned strains, having inherent tolerance of low pH, are suitable for industrial ethanol production.

Ethanol production by *S. cerevisiae* is reported using as substrate a variety of agro-industrial residues, such as enriched pasteurized grape musts, blends of molasses and olive mill wastewaters, olive mill wastewater enriched with glucose, sugarcane bagasse, dried fruits etc., reporting yields up to 0.5 g/g (Behera et al., 2011; Sarris et al., 2014, 2013, 2009; Singh et al., 2013). Other yeasts such as *C. shehatea*, *S. stipitis* and *Pachysolen tannophilus* have been reported as ethanol producers from xylose or cellobiose. Given the fact that *S. cerevisiae* cannot utilize xylose as part of its natural metabolism, the concurrent use of more than one microorganism, for instance, *S. stipitis* and *S. cerevisiae*, has gained popularity in the context of lignocellulose valorization (De Bari et al., 2013; Karagoz et al., 2019; Ntaikou et al., 2018; Santosh et al., 2017).

### 3.5 Single cell oil production

When undiluted PR extract was used as a culture medium for *L. lipofer*, lipid accumulation (L/x%, w/w) was 7.0%, probably restricted by the presence of phenolics or other inhibitors, while when the extract was diluted by 50%, L/x increased to 13.5% (Table 2). Dien et al. (2016) reported that the same strain of *L. lipofer* cultivated in sugar-rich media (i.e. 100 g/L) accumulated 61.6% lipids in the dry cell mass cultivated on glucose, 54.8% on xylose and 56.9% on arabinose. Therefore the low lipid accumulation reported in the current paper should be attributed, in addition to the presence of phenolics, to the low sugar content in the growth medium. On the other hand, yeast growth seems to be unaffected by the concentration of phenolics, as biomass production was proportional to the initial sugar concentration. The biotechnological potential of *M. guilliermondii*, *S. coipomoensis*, and *S. paludigena* to convert lignocellulosic sugars to oily biomass was first reported by Valdés et al. (2020b). In the present study, the above-mentioned strains, cultivated in undiluted PR extract presented an oleaginous capacity depending on the growth conditions (Table 2). Specifically, *M. guilliermondii* was able to produce 2.1 g/L dry cell mass and accumulate 18.1% lipids, while increased lipid accumulation was achieved by *S. paludigena* cultivated under aerobic conditions compared to anaerobic conditions (i.e. 15.4 and 9.0% respectively). Low lipid accumulation was achieved by *S. coipomoensis* under both aerobic and anaerobic conditions (i.e. L/x% = 8.3 and 4.2%), although this yeast was able to accumulate almost 18% of lipids cultivated on glucose and xylose, and 24% on mannose (Valdés et al., 2020b). Although *Y. lipolytica* can grow on a variety of substrates and accumulate lipid in high percentages (Dourou et al., 2018; Papanikolaou and Aggelis, 2011a, 2011b, 2002), during submerged cultures on undiluted PR extract, X_max_ did not exceed 1.7 g/L containing only 7.0% of lipids. However, when PR extract diluted by 50% lipid accumulation increased to 11.0% in the dry cell mass.

Oleic acid (C18:1) was the major FA in all lipids produced by the yeasts, followed by palmitic acid (C16:0) in the lipids of *M. guilliermondii*, *S. coipomoensis*, and *S. paludigena* (Table 4). In the lipids produced by *Y. lipolytica*, C16:0, palmitoleic (C16:1), and linoleic acid (C18:2) participated in non-negligible percentages. Stearic acid (C18:0) was detected in limited amounts in all lipids. According to these data, the oil produced seems to be suitable as raw material in the biodiesel industry. *Y. lipolytica* presented a similar FA profile to that previously reported when cultivated on olive mill wastewater or glycerol (Dourou et al., 2016; Makri et al., 2010; Papanikolaou and Aggelis, 2002). Contrary to the present study, Valdés et al. (2020b) reported that lipids of *M. guilliermondii* and *S. coipomensis* were rich in C16:1, and lipids of *S. paludigena* were rich in C18:2, suggesting that the nature of the carbon source affects the FA profile of these strains.

**Table 4:** Fatty acid composition (%, w/w) of total lipids produced by oleaginous yeasts cultivated on PRs. Data represent the mean of two determinations. Abbreviations: ND, not detected.

| Yeast strain | Fatty acid composition (% , w/w) |  |  |  |  |  |
| --- | --- | --- | --- | --- | --- | --- |
|  | C16:0 | C16:1 | C18:0 | C18:1<br>n-9 | C18:2<br>n-6 | Others |
| *M. guilliermondii* | 35.1<br>± 0.1 | ND | ND | 61.9<br>± 0.9 | ND | 3.0<br>± 0.2 |
| *S. coipomoensis* | 35.5<br>± 0.3 | ND | ND | 62.5<br>± 1.0 | ND | 2.0<br>± 0.9 |
| *S. paludigena* | 26.0<br>± 0.2 | 11.6<br>± 0.3 | 3.9<br>± 0.0 | 37.5<br>± 0.5 | 4.5<br>± 0.0 | 16.7<br>± 1.1 |
| *Y. lipolytica* | 12.7<br>± 0.1 | 14.3<br>± 0.2 | 4.5<br>± 0.1 | 43.2<br>± 0.3 | 19.6<br>± 0.2 | 1.7<br>± 0.5 |

SSF, a fermentation technique suitable for low moisture solid substrates, is frequently used as an effective technology, alternative to submerged fermentation, for the valorization of various agro-industrial by-products (Čertík et al., 2012). PUFA-rich fermented substrates derived from SSFs, after the necessary processing, can be utilized in human diet or as a feed supplement (Čertík et al., 2013; Economou et al., 2010; Gema et al., 2002; Slaný et al., 2021).

Though the presence of R.S. in PRs was high (up to 160 mg/g of substrate, Fig. 3A), *C. echinulata* presented a low ability to grow on PRs used as sole substrate in SSF. This is probably the result of the presence of phenolic compounds in high amounts (i.e. up to 14.0 mg/g substrate, Fig. 3B) or other inhibitors, the high substrate humidity (Fig. 3C) and/or the loss of nutrients during sterilization process. To reduce moisture to adequate levels, blends of PRs with two cereal substrates, WB and OFs, were used. These cereals, treated also as wastes, are a good source of carbon and other nutrients, supporting fungal proliferation (Čertík et al., 2013; Slaný et al., 2020). Actually, their incorporation in high percentage, positively affected fungal growth as R.S. consumption (Fig. 3A) and % substrate utilization (Fig. 3C) confirm. Specifically, when cereals were present at 70 to 90%, % substrate utilization was satisfactory. On the contrary, when PRs were present at a level higher than 30%, R.S. consumption and % substrate utilization were not satisfactory. During mixed cultures, R.S. originated from PRs (0 day in Fig. 3A) were partly utilized by the fungus. Moreover, the addition of WB and OFs resulted in a reduction of water content (Fig. 3C), as well as, of phenolic compounds (up to 3.5 mg/g substrate, Fig. 3B). In all cases, the percentage of substrate utilization and the level of GLA in intracellular lipid gradually increased and maximized on the 5_th_ day (Fig. 3C and Fig. 4). *C. echinulata* cultivated on blends of 90% WB and 10% PRs, produced the highest GLA quantities (i.e. 4.8 mg GLA/g fermented substrate) corresponding to 13.6% GLA in total lipids, while increasing PRs incorporation resulted in decreased GLA production. The addition of 80% OFs resulted in the higher GLA production among OFs-blends (i.e. 3.1 mg GLA/g of fermented substrate representing 4.3% in total lipids). As in the case of WB, when PRs used in a higher percentage (up to 50%) *C. echinulata* formed a low amount of GLA (i.e. 0.5-0.8 mg/g). When PRs were incorporated in low percentages GLA produced was higher than that reported for *C. echinulata* ATHUM 4411 cultivated on orange peel, where GLA production was1.2-1.5 mg/g fermented substrate (Gema et al., 2002).

**Fig. 3:**
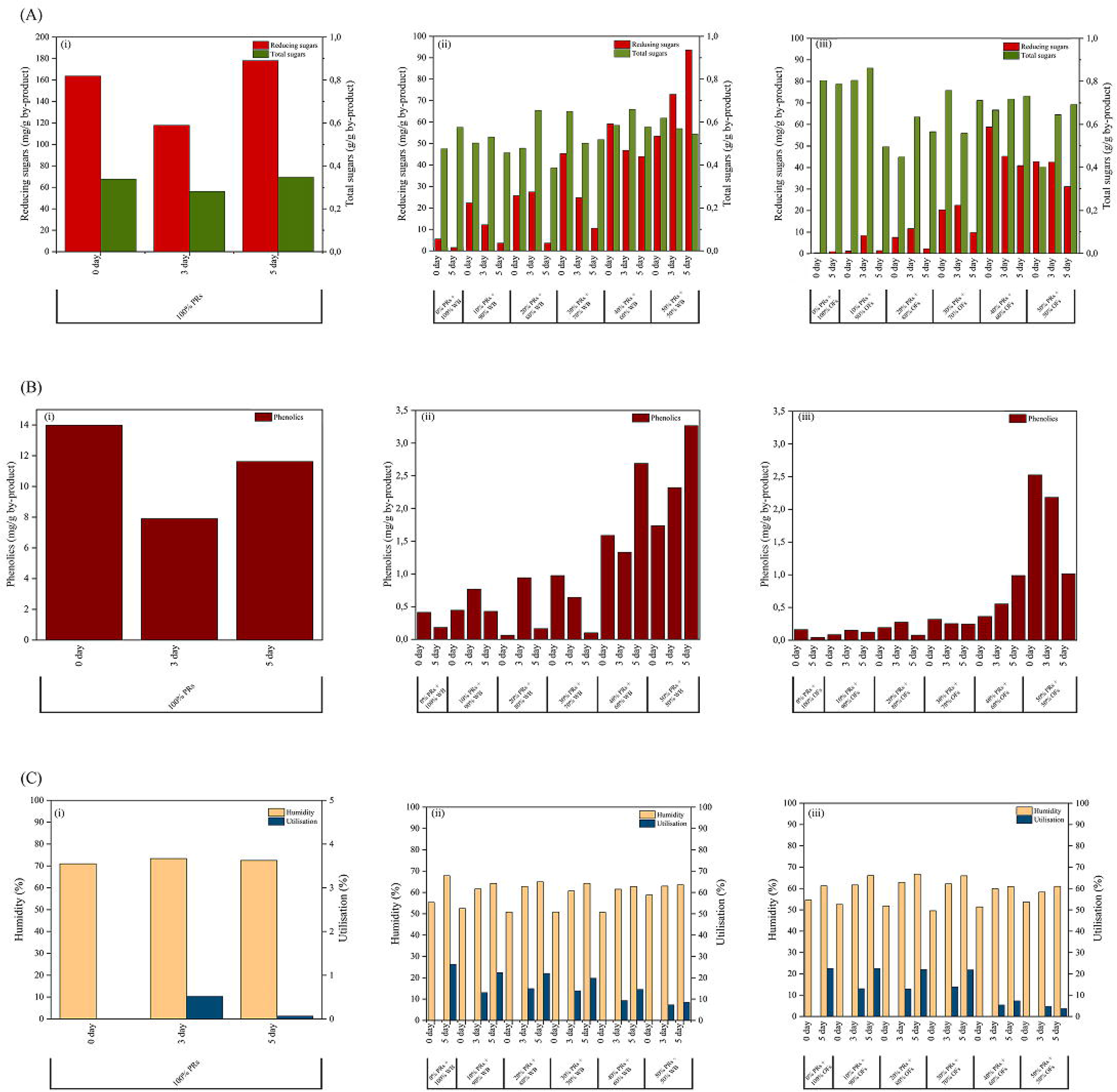
Content of reducing (mg/g by-product) and total sugars (g/g by-product) (A), phenolic compounds (mg/g by-product) (B), and substrate humidity and utilization (%) (C) after solid-state fermentation using *Cunninghamella echinulata* on sole PRs (i), different amounts of PRs and WB (ii), and different amounts of PRs and OFs (iii).

**Fig. 4:**
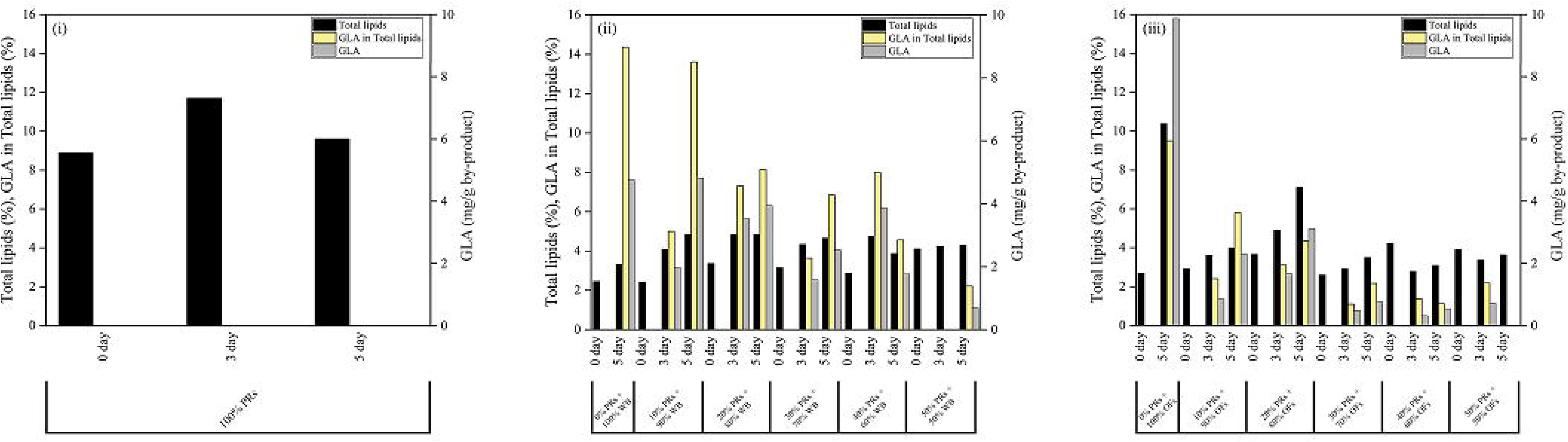
Content of total lipids (%), GLA on total lipids (%), yield of GLA (mg/g by-product) after solid-state fermentation using *Cunninghamella echinulata* on sole PRs (i), different amounts of PRs and WB (ii), and different amounts of PRs and OFs (iii).

The FA composition of lipids extracted from the fermented substrate was similar regardless of the starting substrate (PR or blends of PRs with WB or OFs) (Tables 5 and 6). C18:2 was the dominant FA and its concentration reduced with time. The percentage of C16:0 in total lipids also decreased with time. On the contrary, the percentage of C18:1 increased with time, mostly in WB-blends, suggesting biosynthesis of this FA by *C. echinulata* and/or selective uptake of other FAs resulting in change of lipid profile. As expected GLA was not detected in the initial substrate and therefore was synthesized by the fungus during substrate assimilation. Some quantities of C18:2 were probably converted to GLA. In all cases, the final fermented substrate containing C18:2, GLA and C18:3n-3, even in small quantities, is of high nutritional value for animal or human consumption. The necessity for cereal-based products enriched with essential FAs and other compounds (such as pigments) has been reported (Čertík et al., 2013; Certik and Adamechova, 2009) and the use of oleaginous fungi during SSF seems like a promising approach.

**Table 5:** Fatty acid composition (%) of total lipids of *Cunninghamella echinulata* during solid-state fermentation on PRs and WB used in different ratios. Data represent the mean of two determinations. Abbreviations: ND, not detected

| Ratio of<br>PRs:WB (%) |  | Day of<br>SSF | Fatty acid composition (%) |  |  |  |  |  |  |  |
| --- | --- | --- | --- | --- | --- | --- | --- | --- | --- | --- |
| PRs | WB |  | C16:0 | C16:1 | C18:0 | C18:1<br>n-9 | C18:2<br>n-6 | C18:3<br>n-6 | C18:3<br>n-3 | Others |
| 0 | 100 | 0 | 19.6<br>± 0.2 | ND | 3.7<br>± 0.3 | 24.5<br>± 0.9 | 48.6<br>± 1.3 | ND | 3.3<br>± 0.2 | 0.3<br>± 0.1 |
|  |  | 5 | 14.6<br>± 0.1 | 0.6<br>± 0.0 | 2.9<br>± 0.1 | 26.4<br>± 1.1 | 39.8<br>± 1.1 | 14.4<br>± 0.4 | 1.1<br>± 0.0 | 0.2<br>± 0.0 |
| 10 | 90 | 0 | 20.1<br>± 0.7 | 0.6<br>± 0.0 | 1.3<br>± 0.0 | 25.2<br>± 1.2 | 48.6<br>± 0.6 | ND | 3.6<br>± 0.1 | 0.3<br>± 0.1 |
|  |  | 3 | 15.7<br>± 0.3 | 0.9<br>± 0.1 | 4.4<br>± 0.5 | 29.7<br>± 0.7 | 40.5<br>± 0.7 | 5.0<br>± 0.1 | 3.3<br>± 0.2 | 0.5<br>± 0.2 |
|  |  | 5 | 13.9<br>± 0.1 | 0.8<br>± 0.2 | 3.5<br>± 0.2 | 32.1<br>± 1.0 | 34.1<br>± 0.3 | 13.6<br>± 0.2 | 1.7<br>± 0.0 | 0.3<br>± 0.0 |
| 20 | 80 | 0 | 19.9<br>± 0.5 | 0.2<br>± 0.0 | 2.2<br>± 0.0 | 26.0<br>± 0.9 | 48.5<br>± 1.2 | ND | 3.0<br>± 0.2 | 0.2<br>± 0.0 |
|  |  | 3 | 16.6<br>± 0.4 | 0.5<br>± 0.0 | 5.5<br>± 0.4 | 30.9<br>± 0.2 | 37.2<br>± 1.0 | 7.3<br>± 0.1 | 1.6<br>± 0.0 | 0.4<br>± 0.1 |
|  |  | 5 | 16.5<br>± 0.5 | 0.8<br>± 0.1 | 5.9<br>± 0.6 | 33.9<br>± 0.7 | 33.3<br>± 0.4 | 8.1<br>± 0.2 | 1.1<br>± 0.1 | 0.4<br>± 0.1 |
| 30 | 70 | 0 | 19.9<br>± 1.0 | 1.0<br>± 0.3 | 2.5<br>± 0.1 | 25.3<br>± 0.3 | 47.6<br>± 0.9 | ND | 3.5<br>± 0.3 | 0.2<br>± 0.0 |
|  |  | 3 | 18.0<br>± 0.2 | 0.4<br>± 0.0 | 4.3<br>± 0.1 | 26.5<br>± 0.5 | 44.2<br>± 0.7 | 3.6<br>± 0.0 | 2.7<br>± 0.2 | 0.3<br>± 0.0 |
|  |  | 5 | 17.6<br>± 0.1 | 1.0<br>± 0.0 | 5.9<br>± 0.3 | 34.5<br>± 1.1 | 32.2<br>± 0.7 | 6.8<br>± 0.5 | 1.6<br>± 0.1 | 0.4<br>± 0.1 |
| 40 | 60 | 0 | 21.2<br>± 1.1 | 0.5<br>± 0.1 | 3.0<br>± 0.2 | 23.8<br>± 0.6 | 48.2<br>± 0.2 | ND | 3.1<br>± 0.2 | 0.2<br>± 0.0 |
|  |  | 3 | 20.2<br>± 0.9 | 0.3<br>± 0.0 | 2.4<br>± 0.2 | 25.1<br>± 0.5 | 47.4<br>± 1.4 | 1.3<br>± 0.0 | 3.0<br>± 0.9 | 0.3<br>± 0.0 |
|  |  | 5 | 17.3<br>± 0.3 | 0.4<br>± 0.0 | 4.3<br>± 0.7 | 30.3<br>± 1.0 | 40.3<br>± 1.1 | 4.6<br>± 0.2 | 2.5<br>± 0.7 | 0.3<br>± 0.1 |
| 50 | 50 | 0 | 19.3<br>± 0.4 | 0.3<br>± 0.1 | 2.1<br>± 0.1 | 23.6<br>± 0.2 | 52.6<br>± 0.9 | ND | 0.2<br>± 0.0 | 1.9<br>± 0.3 |
|  |  | 3 | 16.2<br>± 0.1 | 0.2<br>± 0.0 | 3.2<br>± 0.0 | 23.3<br>± 0.5 | 51.7<br>± 0.6 | 5.0<br>± 0.1 | 0.2<br>± 0.0 | 0.2<br>± 0.0 |
|  |  | 5 | 17.9<br>± 0.3 | 0.8<br>± 0.2 | 3.4<br>± 0.2 | 24.6<br>± 0.4 | 46.1<br>± 1.0 | 2.2<br>± 0.1 | 4.1<br>± 0.2 | 0.9<br>± 0.2 |

**Table 6:**
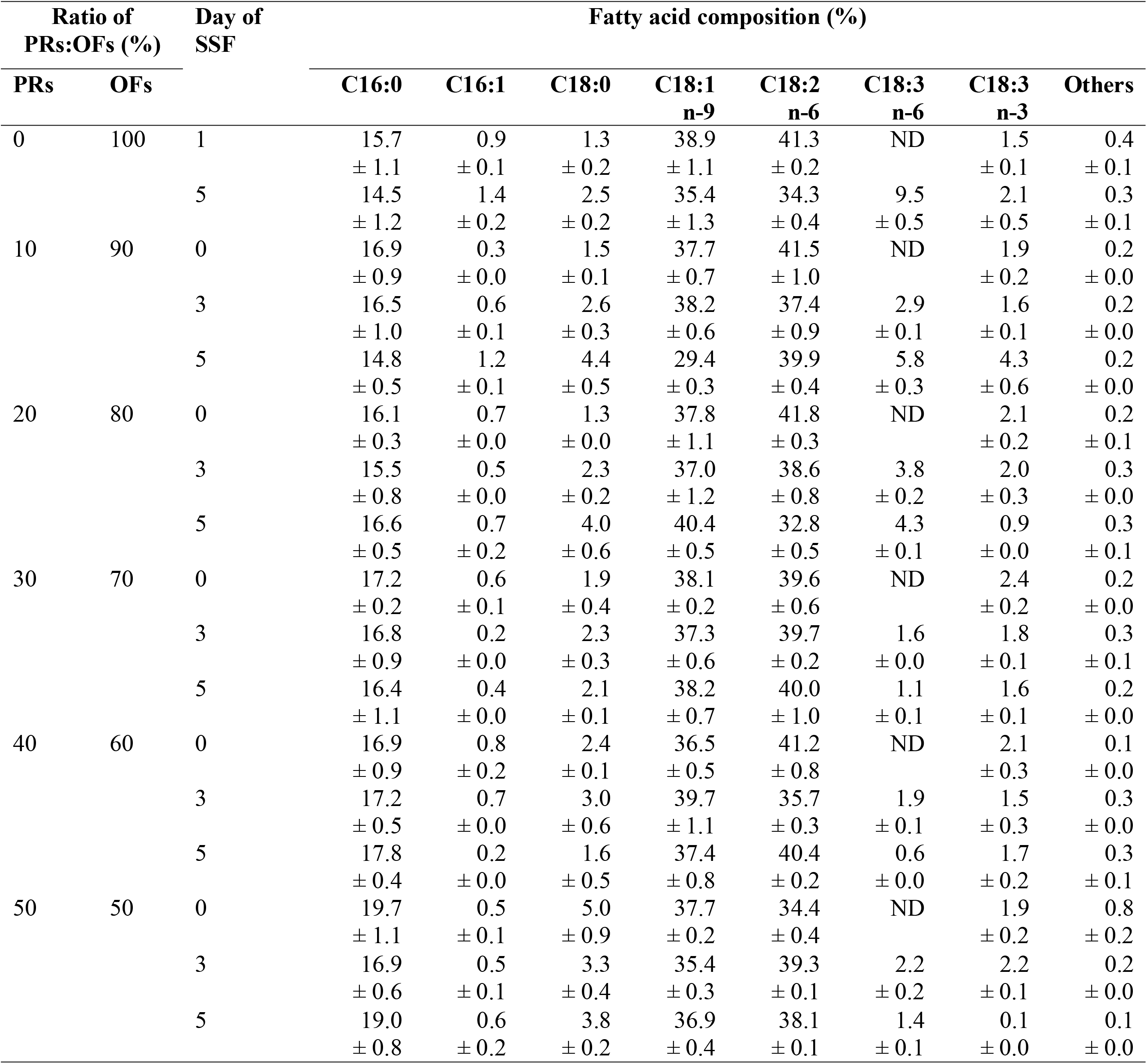
Fatty acid composition (%) of total lipids of *Cunninghamella echinulata* during solid-state fermentation on PRs and OFs used in different ratios. Data represent the mean of two determinations.

TLC analysis of total lipids by *C. echinulata* is depicted in Fig. 5. The lipids of the fermented PRs contained various species of TAGs, while the absence of 1,3-DAG and ergosterol is noticed. Lipids from SSF performed on WB and OF alone contained ergosterol in significant percentages and TAGs were present in greater diversity. As the incorporation of PRs increased in the substrate, specific species of TAGs (i.e. TAG1) also increased, while on the contrary, the ergosterol content decreased. There are several distinct patterns of sterols among fungi, depending on the phylum and/or environmental conditions (Volkman, 2003). The presence of ergosterol in fungal or yeast lipids is essential for their response to stress by inhibitors and therefore to their growth (Dupont et al., 2012). Phytosterol content was similar in lipids produced during SSF on individual substrates (approximately 11%, w/w) and decreased in blends of PRs with cereals, with exception of the substrate in which OFs participated at a percentage of 80%, in which phytosterol remained in high levels. Moreover, the incorporation of WB resulted in a higher amount of free FAs on the final product, while the percentage of polar lipids slightly increased in the case of blends of PRs with cereals (ranging from 6.2 to 11.5%). During SSF of WB by *Umbelopsis isabellina* the percentage of free FAs and sterols increased (Slaný et al., 2021).

**Fig. 5:**
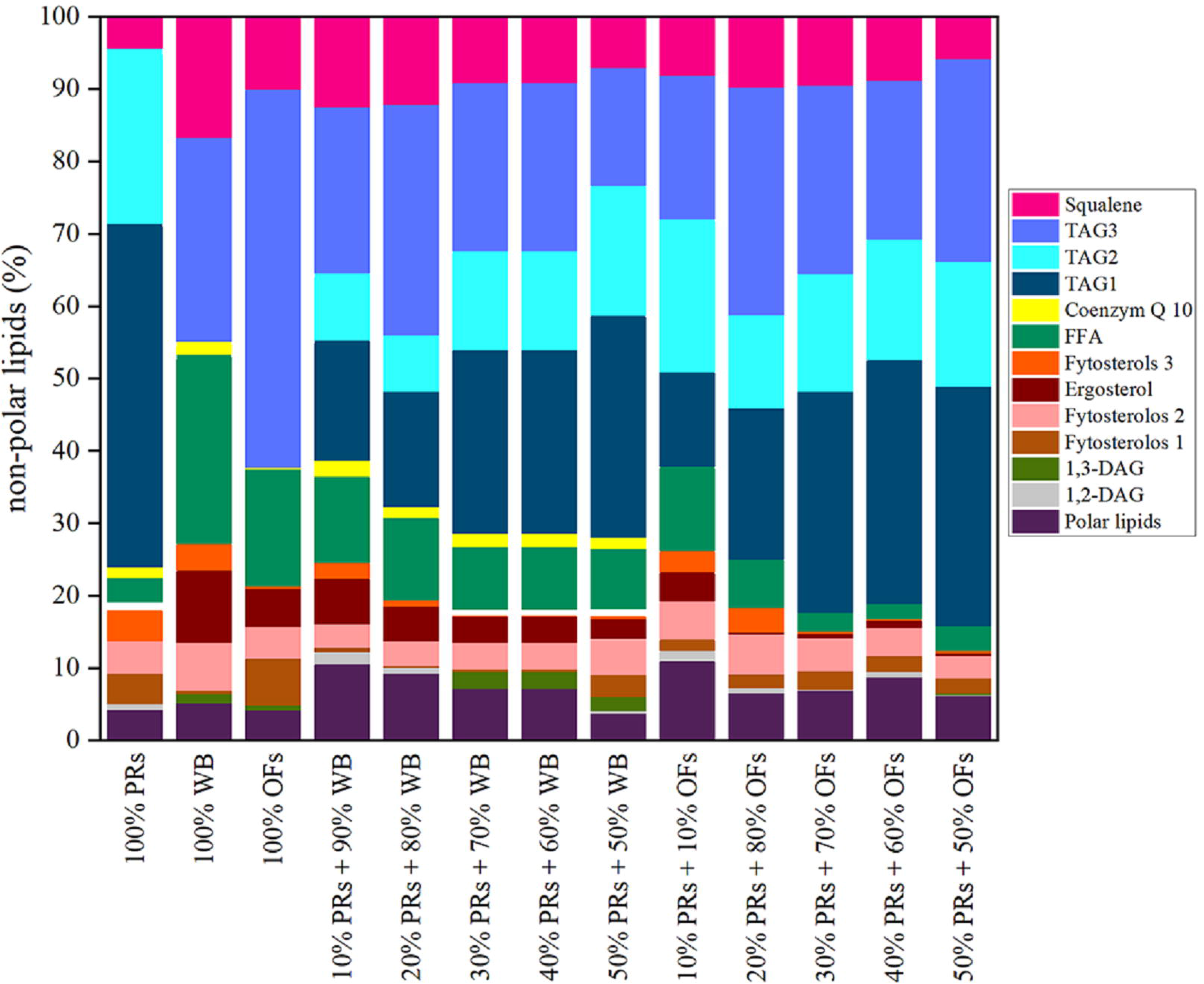
TLC analysis of non-polar lipids of *Cunninghamella echinulata*.

## 4. Conclusions

Agro-industrial residues, such as PRs, are suitable low-cost substrates for microbial fermentation processes leading to a “green” and cost-effective production of value-added products. These residues are largely available and abundant in nutrients. Researchers are focused on pretreatment methods of lignocellulosic biomass, while they have to overcome a variety of peculiarities to use such materials. In this investigation, efficient extraction and hydrolysis of the polysaccharides found in PRs were conducted in a one-step process and the resulted broth was successfully utilized by several yeast strains, able to grow into such extreme conditions, for either ethanol or SCOs production. Moreover, PRs combined with cereals, were effectively operated by an oleaginous fungus for the production of a final fermented substrate enriched with PUFAs. We conclude that new perspectives are emerging for the use of PRs as a raw material to produce valuable microbial metabolites, using environmentally friendly approaches.

## Abbreviations

FA: fatty acid
GLA: γ-linolenic acid
OFs: oat flakes
PRs: pomegranate residues
PUFAs: polyunsaturated fatty acids
R.S.: reducing sugars
SCOs: single cell oil
SSF: Solid-state fermentation
TAGs: triacylglycerols
T.S.: total sugars
WB: wheat bran

